# Silencing hippocampal CA2 reduces behavioral flexibility in spatial learning

**DOI:** 10.1101/2022.11.08.515580

**Authors:** Andrew B. Lehr, Frederick L. Hitti, Scott H. Deibel, Tristan M. Stöber

## Abstract

The hippocampus is a key structure involved in learning and remembering spatial information. However the extent to which hippocampal region CA2 is involved in these processes remains unclear. Here we show that chronically silencing dorsal CA2 impairs reversal learning in the Morris water maze. After platform relocation, CA2-silenced mice spent more time in the vicinity of the old platform location and less time in the new target quadrant. Accordingly, behavioral strategy analysis revealed increased perseverance in navigating to the old location during the first day and an increased use of non-spatial strategies during the second day of reversal learning. Confirming previous indirect indications, these results demonstrate that CA2 is recruited when mice must flexibly adapt their behavior as task contingencies change. We discuss how these findings can be explained by recent theories of CA2 function and outline testable predictions to understand the underlying neural mechanisms. Demonstrating a direct involvement of CA2 in spatial learning, this work lends further support to the notion that CA2 plays a fundamental role in hippocampal information processing.

## 1 Introduction

The hippocampus is crucially involved in spatial memory processing. While the dentate gyrus (Sutherland et al., 1983), CA3 (Nakazawa et al., 2003), and CA1 (Tsien et al., 1996) are important for spatial learning and memory, existing experimental work suggests that CA2 may not be required. In the Morris water maze, a gold standard test for hippocampus-dependent spatial learning (Morris et al., 1982; Morris, 1984), mice with chronically silenced CA2 learned to find a hidden platform over the course of five training days during both initial learning and reversal learning (Hitti and Siegelbaum, 2014). While animals could learn the task, Hitti and Siegelbaum (2014) noted a trend towards CA2-silenced mice learning more slowly, potentially hinting at a subtle impairment.

Understanding why subtle differences in CA2-silenced mice may arise in the Morris water task requires a clear definition of the neural computations that must be performed. To directly navigate to the submerged platform the animal must form an internal representation of the location based on its behavioral experience. Upon platform relocation during reversal learning, an additional spatial representation is required and the animal must learn that the novel location is now rewarding but the previous location no longer is. To perform such an update, the animal must first detect that task contingencies have changed before it can use the new memory traces to update its internal representations. This means that while initial learning requires forming a new representation, reversal learning additionally requires detecting the change and prioritizing new vs. old memory traces.

Recent experimental studies have shown that CA2 is indeed recruited when new representations need to be formed, suggesting it may be involved both in learning and reversal learning during the Morris water task. Introducing inanimate objects or conspecifics in otherwise identical environments leads to the recruitment of new cell assemblies (Wintzer et al., 2014), spatial remapping (Alexander et al., 2016) and the encoding of such information in CA2 population activity (Donegan et al., 2020; Boyle et al., 2022; Oliva et al., 2020), with larger effects compared to other hippocampal subregions. Accordingly, animals with chronically silenced CA2 show slower contextual habituation, indicating a role in forming representations during spatial learning (Boehringer et al., 2017).

CA2 may be even more important for reversal learning. When local cues are rotated, akin to the platform relocation during reversal learning (180^*°*^ rotation), CA2 place fields rotate with them (Lee et al., 2015). Further, during reversal learning new memory traces should be prioritized over old ones to update internal representations. This requires consolidation of the new experience into memory, a process known to depend crucially on sharp wave ripples during which memory traces are reactivated (Girardeau et al., 2009; Jadhav et al., 2012; Ego-Stengel and Wilson, 2010; Oliva et al., 2020). Reactivation quality has been shown to predict task performance (Dupret et al., 2010), especially during early learning (Singer et al., 2013). When CA2 is silenced, reactivation quality deteriorates, and multiple different experiences no longer remain properly separated. He et al. (2021) showed that in control animals single memory traces tend to be reactivated individually during single sharp wave ripples. In contrast, transient silencing of CA2 increased simultaneous reactivation of memory traces relating to two separate spatial environments. For reversal learning, these sharp wave ripple related changes may be detrimental, as poorer replay quality and simultaneous reactivation of both the old and new memory traces would make it difficult for CA2-silenced animals to update internal representations and flexibly adapt their behavior.

Based on these findings, we hypothesized that CA2 silencing should negatively affect both early learning and reversal learning, when new representations needs to be formed. However, the detrimental effect should be stronger during reversal learning, where in addition old memory traces may interfere with the reactivation of new traces. This motivated us to further analyze the Morris water maze data from Hitti and Siegelbaum (2014). Employing a broad set of analysis tools, we searched for a specific impairment of mice with chronically silenced dorsal CA2 during early learning phases. While we could not detect consistent differences during initial learning, we found profound effects during early reversal learning. CA2-silenced mice spent more time near the old platform location, perseverated more strongly during the first day and employed more non-spatial search strategies during the second day of reversal learning. Taken together, these results indicate hippocampal region CA2 contributes to flexibly adapting behaviour upon changes to the local environment.

## 2 Methods

### 2.1 Animals

Animals were housed in groups of two to five per cage with *ad-libitum* supply of food and water. Synaptic output of CA2 pyramidal cells was blocked by selectively expressing tetanus neurotoxin light chain (TeNT) in these cells. A Cre-dependent adeno-associated virus vector carrying eGFP–TeNT was bilaterally injected in the dorsal hippocampus of Amigo2-Cre+ mice (*n*_*t*_ = 10, all male), which selectively express Cre in CA2 pyramidal cells. Control mice (*n*_*c*_ = 8, all male) received the same treatment but the viral vector carried only yellow fluorescent protein (YFP) instead of TeNT. In this study we further analyzed data from a published experiment approved by the Columbia University Institutional Animal Care and Use Committee. For an exhaustive description of animal treatment, please refer to the original publication (Hitti and Siegelbaum, 2014).

### 2.2 Morris water maze task

To test spatial memory, mice went through a fixed 14 day test regime in the Morris water maze. These tests were conducted during the light phase of a 12h light-dark cycle and after three days habituation to handling and transport. Before each session mice were allowed to habituate for 1h. Mice had to find a submerged platform in a pool with a diameter of 120 cm containing opacified water. Animal position was tracked with a FireWire camera and ANY-maze recording software. Each day consisted of four 1 minute trials with an intertrial interval of around 20 minutes and for each trial animals were placed at a random start position. During day 1 and 2 the platform was marked with a flag and distal cues were concealed. The platform was moved to a new location for each trial. If an animal failed to find the platform within 1 minute, it was guided to the platform. On day 3 to 7 the flag was removed from the platform, curtains hiding distal cues were removed, and the platform remained in the same location in the southwest quadrant for all acquisition trials. On day 8, a single probe trial was conducted in which the platform was removed and animals swam for one minute. For reversal training, the platform was moved to the northeast quadrant and the procedure repeated. A 1 minute probe trial on day 14 again without the hidden platform marked the end of the experiment for each animal. The experimenter had no information about the treatment groups.

### 2.3 Data analysis

Experimental data and the complete analysis is publicly available at https://github.com/tristanstoeber/CA2_spatiallearning.

#### 2.3.1 Preprocessing

Before the analysis, invalid values were excluded and spatial coordinates linearly interpolated with a fixed temporal resolution of Δ*t* = 0.1*s*.

#### 2.3.2 Occupancy maps

To create occupancy maps for each animal we binned spatial coordinates with Δ*x* = 0.025*m*, summed the time spent in each bin across the four trials from that day, and smoothed with a Gaussian filter with *σ* = 0.025*m*. Finally, we averaged over the occupancy map across animals in each treatment group.

#### 2.3.3 Mean distance to platform

To calculate the mean distance to the learning and relearning platforms we computed the arithmetic mean over the distance between the center of the animal, as determined by the ANY-maze recording software, and the center of the platform at each timepoint. We tested for statistical differences between treatment groups by performing two-way repeated measures ANOVA separately for the learning, day 3 to 7, as well the reversal learning phase, day 9 to 13 with the rstatix package in R (R Core Team, 2021).

#### 2.3.4 Fraction of time spent in target quadrant

To determine the fraction of time spent in each quadrant we used the xy-coordinates from the ANY-maze recording software. After counting the time spent in each quadrant, we divided by the total trial duration. ANOVA tests were performed analog to the mean distance to platform.

#### 2.3.5 Strategy analysis

To infer navigational strategies we used the Rtrack package (Overall et al., 2020). Since its based on a parameter-free machine learning approach, no custom parameters were required. We compared perseverance behaviour on day 9 and with a one-tailed Mann-Whitney U test with the alternative hypothesis that the fraction of trials with perseverance for CA2-silenced mice is greater compared to control mice. We compared non-spatial strategies on each day of learning and reversal learning with a one-tailed Mann-Whitney U test with the alternative hypothesis that the fraction of trials with non-spatial strategies is greater in CA2-silenced mice during early learning and early reversal learning.

## 3 Results

### 3.1 CA2-silenced mice spend more time near the old platform location during reversal learning in the Morris water maze

To test our hypothesis that CA2 silencing affects the early stages of learning and reversal learning, we decided to analyze Morris water maze data from Hitti and Siegelbaum (2014). In each phase CA2-silenced (*n*_*t*_ = 10) and control (*n*_*c*_ = 8) mice were trained for five days with four trials per day to locate a hidden platform, submerged in opacified water. Mice with chronically silenced dorsal CA2 successfully learned to navigate to the platform in both phases. Hitti and Siegelbaum (2014) showed that during probe trials, in which the platform was removed, CA2-silenced animals spent the same amount of time in the respective target quadrant as control mice. Further, they did not find a significant difference in performance during acquisition as quantified by the distance travelled and time to reach the platform across acquisition trials. However, the authors reported a trend towards longer path lengths and latencies to reach the hidden platform for CA2-silenced mice.

Given our hypothesis that early spatial learning should be impaired in CA2-silenced mice, we performed post-hoc tests for each day of learning and reversal learning. The analysis revealed that CA2-silenced mice indeed required longer path lengths and trial durations to reach the hidden platform on the second day of learning (day 4: path length *p* = 0.04, trial duration *p* = 0.03, Mann-Whitney U test, one-tailed, *n*_*t*_ = 10, *n*_*c*_ = 8, no correction for multiple comparison) and on the second and last day of reversal learning (day 10: path length *p* = 0.005, trial duration *p* = 0.006; day 13: path length *p* = 0.02, trial duration *p* = 0.04; Mann-Whitney U test, one-tailed, *n*_*t*_ = 10, *n*_*c*_ = 8, no correction for multiple comparison). This agrees with Hitti and Siegelbaum’s observation of a trend towards slower acquisition in CA2-silenced mice, which is visible on the second day of learning (day 4) and the second day of reversal learning (day 10) (compare Supplementary Fig. 1 as well as Extended Data Fig. 6b,c in Hitti and Siegelbaum, 2014). Notably the post-hoc tests demonstrate that the effect was most robust for early reversal learning.

To visualize potential differences, next we plotted occupancy maps averaged over all animals for each day for learning and reversal learning. While it is difficult to see a systematic difference in learning (see Supplementary Fig. 2), occupancy maps during reversal learning vividly show that CA2-silenced animals spend more time in the proximity of the old platform location (Fig. 1D). This suggests that CA2-silenced animals may indeed have impairments in adapting their behavioral response after the platform location is changed, but not necessarily during initial learning.

To quantify this effect, we measured the mean distance to the original platform location over the course of learning and reversal learning. Both CA2-silenced and control animals performed similarly during learning (two-way repeated-measures ANOVA: *F* (1, 16) = 0.10, *p* = 0.75, *n*_*t*_ = 10, *n*_*c*_ = 8), supporting the visual impression from the occupancy maps. For reversal learning, if CA2-silenced animals had difficulties shifting their response, we would expect them to be closer on average to the original platform location specifically after platform reversal. Indeed during reversal learning, the mean distance to the old platform location was lower for CA2-silenced animals (see Fig. 1B, two-way repeated-measures ANOVA: *F* (1, 16) = 6.53, *p* = 0.02, *n*_*t*_ = 10, *n*_*c*_ = 8) and there was a consistent trend towards more distance to the new platform location (see Supplementary Fig. 3, two-way repeated-measures ANOVA: *F* (1, 16) = 2.92, *p* = 0.11, *n*_*t*_ = 10, *n*_*c*_ = 8). Accordingly, the fraction of time CA2-silenced animals spent in the target quadrant was lower than controls during reversal learning (see Fig. 1C, two-way repeated-measures ANOVA: *F* (1, 16) = 4.87, *p* = 0.04, *n*_*t*_ = 10, *n*_*c*_ = 8), while during learning both groups performed the same (two-way repeated-measures ANOVA: *F* (1, 16) = 0.15, *p* = 0.71, *n*_*t*_ = 10, *n*_*c*_ = 8)). Taken together, CA2-silenced animals did not show reliable differences to controls during the initial learning phase of the Morris water task, however they differed from controls during reversal learning, where they spent more time around the old platform location.

### 3.2 CA2-silenced mice show increased perseverance at the old location and delayed onset of spatial search strategies for the new location during early reversal learning

Based on previous published theoretical arguments (Stöber et al., 2020, see discussion), we hypothesized that CA2-silenced mice should be impaired in changing their behavioural strategy upon changing task contingencies. In particular, we expected that CA2-silenced animals show increased perseverance to navigate to the old platform location directly after platform reversal. Thus, we classified navigation strategies with *Rtrack*, a parameter free random forest classifier trained on human curated example data (Overall et al., 2020; Berdugo-Vega et al., 2020, 2021). We distinguished each trial into non-spatial and spatial strategies – *thigmotaxis, circling, random path, scanning* as well as *chaining, directed search, corrected search, direct path*, and *perseverance* (see Fig. 2B).

**Figure 1:**
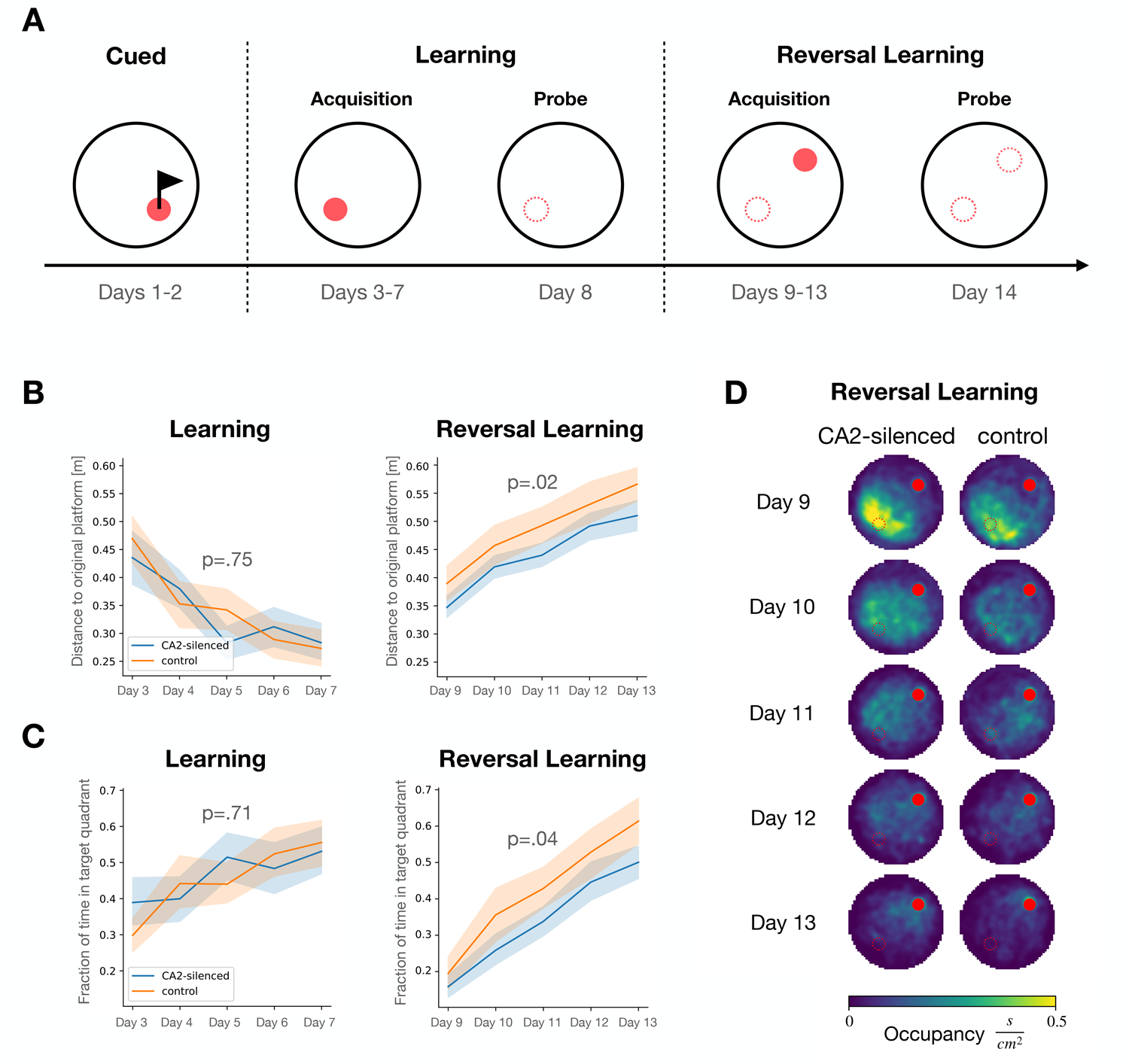
CA2-silenced mice spend more time near the old platform location during reversal learning in the Morris water maze. **(A)** Experimental procedure consisted of three phases: cued, learning, and reversal learning. During the cued phase the platform was cued with a flag and relocated for each trial. In subsequent phases the platform was hidden. Learning consisted of five days for acquisition with four trials per day and a sixth day with a single probe trial. The hidden platform was moved on Day 9 and this procedure was repeated. **(B)** During reversal learning CA2-silenced (blue) were consistently closer to the original platform compared to control mice (blue), as measured by the average distance to the original platform. **(C)** During reversal learning the fraction of time spent in the target quadrant was consistently lower for CA2-silenced (blue) compared to control mice (orange). During the learning phase, neither the average distance to the platform location (B), nor the fraction of time spent in the target quadrant (C) differed between the groups. In both panels (B) and (C) the line shows the mean and shaded regions the 95% confidence interval. **(D)** Average occupancy maps show that CA2-silenced mice spent more time around the old platform location compared to control mice during reversal learning (Days 9-13). Yellow represents high and blue low occupancy on a linear scale. Red filled circle demarcates current platform location and unfilled circles previous locations.

**Figure 2:**
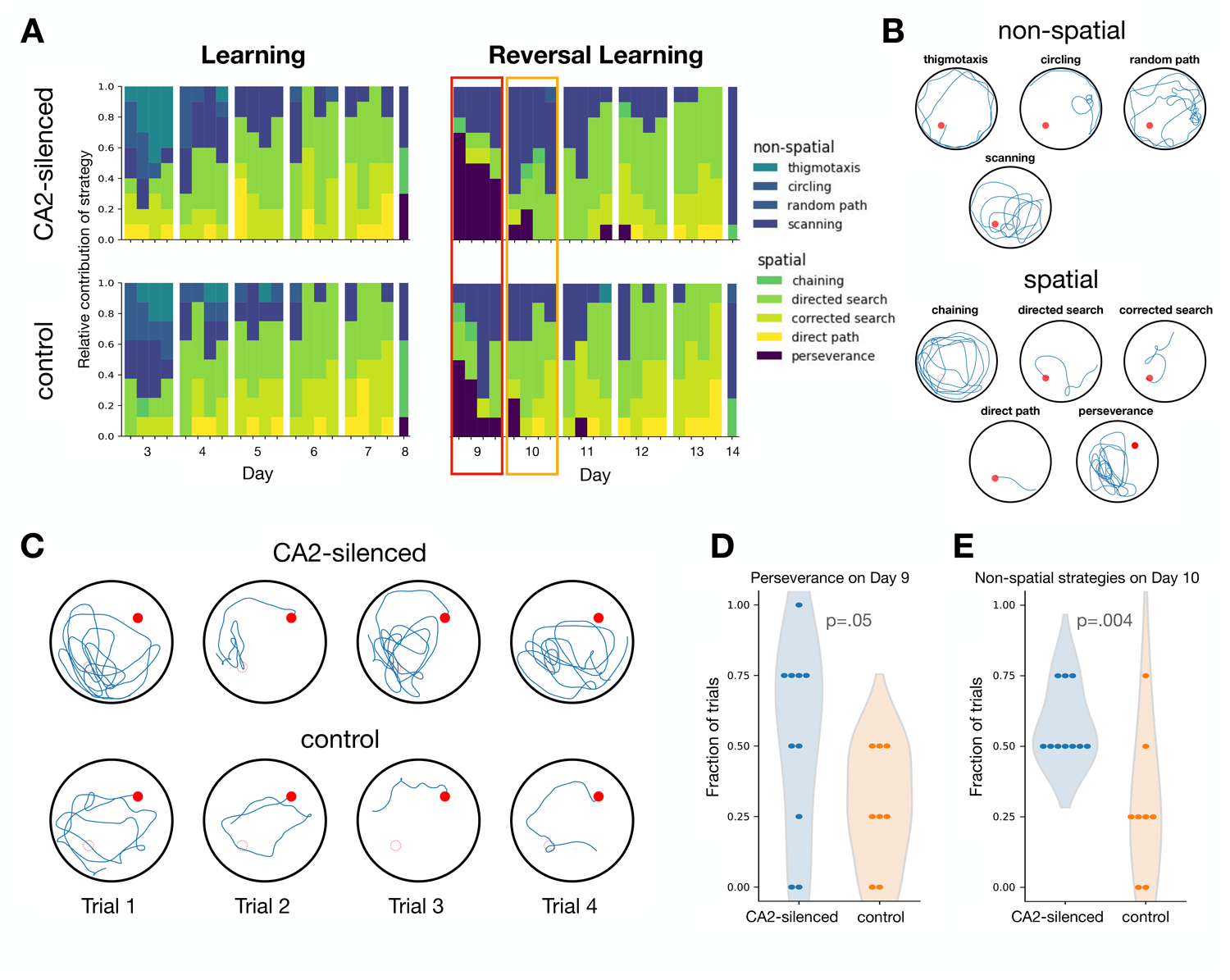
CA2-silenced mice show increased perseverance in navigating to the old location during the first day and an increased use of non-spatial strategies during the second day of reversal learning. **(A)** Relative contribution of behavioral strategies are shown across the learning and reversal learning phases for CA2-silenced (top) and control (bottom) animals. Red box around Day 9 highlights increased perseverance and orange box around Day 10 highlights delayed onset of spatial strategies in CA2-silenced animals after platform reversal (compare panel D and E respectively). **(B)** Examples of behavioral strategies separated into non-spatial and spatial as classified by Rtrack. **(C)** Example paths from Day 9 for a CA2-silenced animal that continued to persevere (top) and a control animal that adapted its strategy (bottom) from scanning to directed/corrected search. **(D)** Fraction of trials classified as perseverance strategy on Day 9. **(E)** Fraction of trials classified as non-spatial strategy on Day 10. In panels (D) and (E) each data point represents one animal. Shaded region is a kernel density estimate of the probability density function.

In accordance with our hypothesis, we observed that CA2-silenced animals tend to persevere in searching near the old platform location on the first day of reversal learning while controls quickly adapt their strategy (Fig. 2A). This is evidenced by a larger fraction of trials on which CA2-silenced animals showed perseverance on this day (Fig. 2D, Day 9: Mann-Whitney U, *p* = 0.05, one-tailed). To exemplify the contrast we plotted the trajectories of two extreme cases, a CA2-silenced mouse that showed perseverance in all four trials and a control mouse with no perseverance (Fig. 2C). While the CA2-silenced mouse continues to persevere at the old platform location, the control mouse switches its strategy over the course of the four trials from scanning to directed/corrected search.

After increased perseverance at the old location on the first day of reversal learning, CA2-silenced animals showed more non-spatial search strategies on the second day of reversal learning Fig. 2E, Mann-Whitney U, *p* = 0.004, one-tailed, *n*_*t*_ = 10, *n*_*c*_ = 8). Otherwise during the remaining days of learning and reversal learning CA2-silenced mice employed a similar proportion of non-spatial strategies to controls (Mann-Whitney U, *p >* 0.19, one-tailed, *n*_*t*_ = 10, *n*_*c*_ = 8, no correction for multiple comparison).

## 4 Discussion

Here we have shown that hippocampal region CA2 contributes to spatial learning in the Morris water maze – in particular to early reversal learning. On the first day of reversal learning CA2-silenced mice expressed more perseverance in navigating to the old platform location and on the second day they employed fewer spatial search strategies. Consistent with this, they also spent less time in the target quadrant and, instead, swam in closer proximity to the old platform location throughout the reversal learning phase. Despite these differences, CA2-silenced animals were still able to learn to find a hidden platform over five days of reversal training in the Morris water maze and over repeated trials successfully adapted their behavior as evidenced by comparable performance to controls in the final probe trial (see Hitti and Siegelbaum, 2014).

In contrast to our initial hypothesis, we were unable to detect consistent differences in the learning phase upon silencing dorsal CA2. It remains to be tested whether instead ventral CA2 may contribute to initial learning. Such a functional differentiation between dorsal and ventral CA2 would reconcile our findings with other indications for a role of CA2 in initial learning (see subsections 4.1 and 4.5).

### 4.1 Our findings corroborate previous indications for CA2’s role in spatial learning

Lee et al. (2010) were the first to establish a link between CA2 and spatial learning and memory in the Morris water maze. They globally knocked out RGS14, which is almost exclusively expressed both in dorsal and ventral CA2 pyramidal cells (see also Dudek et al., this issue). Knocking out RGS14 unlocked otherwise restricted plasticity (Zhao et al., 2007; Chevaleyre and Siegelbaum, 2010) at CA3→CA2 synapses and accelerated initial learning in the Morris water maze, with no difference during the probe trial. It remains to be tested whether the same manipulation increases or deteriorates performance during reversal learning.

Other indirect evidence comes from the inbred BTBR mouse strain with an autism-like phenotype (Moy et al., 2007). Besides impairments in sociability and social memory, BTBR mice require more time to learn the Morris water maze and show no significant quadrant preference during the reversal probe trial (Moy et al., 2007). Interestingly, several of the anatomical and physiological aberrations of BTBR mice (Dodero et al., 2013; Ellegood and Crawley, 2015; Gozzi and Schwarz, 2016) are CA2 specific (Cope et al., 2021): (i) fewer mossy fibers from adult born granule cells to CA2, (ii) smaller CA2 volume, (iii) fewer dendritic spines, microglia and astroctyes and (iv) stronger expression of perineuronal nets in CA2. Restoring levels of perineuronal nets in BTBR mice partially recovers social recognition memory, demonstrating an involvement of CA2 in the BTBR behavioral phenotype (Cope et al., 2021). However, whether the deficit in spatial learning of BTBR mice depends on changes to CA2 remains to be investigated.

Another indication for an involvement of CA2 in spatial learning comes from a knockout study of the mineralocorticoid receptor (Berger et al., 2006). Within the hippocampus the mineralocorticoid repector is most strongly expressed in CA2 pyramidal cells (Kretz et al., 2001; McCann et al., 2019). Without mineralocorticoid receptor expression the molecularly distinct nature of CA2 seems to disappear (McCann et al., 2019): CA2 specific markers, such as NECAB2, PCP4 and WFA are reduced, and otherwise restricted CA3→CA2 plasticity can be readily induced. Knocking out mineralocorticoid receptors in mouse forebrain, which includes the hippocampus and presumably strongly impacts CA2 given its high receptor expression, leads to deficiencies in the Morris water maze (Berger et al., 2006). Mice are slower in learning the initial platform location and upon platform relocation some animals show strong perseverative searching at the old platform location (Berger et al., 2006).

In each of these studies, CA2 appears to play a role in initial learning, and if tested, also in reversal learning. In contrast to our study, the described hippocampal manipulations appear to be global, involving both dorsal and ventral CA2. This may resolve the initial learning differences compared to our study (for further discussion see subsection 4.5).

Beyond behavioral evidence, recent work has established a link between CA2 and the formation of spatial maps within the hippocampus. Transiently silencing CA2 pyramidal cells slowed emergence and reduced trial-to-trial stability of CA1 place fields (He et al., 2022). The same manipulation also reduced the synchrony between hippocampal subregions during sharp wave-ripples. It is conceivable that such impairments may reduce performance in the Morris water maze.

### 4.2 Recent theories of CA2 function are consistent with reduced flexibility in CA2-silenced mice: sensory-memory conflicts and prioritization for replay

Perseverance at the old platform location in CA2-silenced animals is consistent with two current theories of CA2 function. Middleton and McHugh (2019) suggested that CA2 is well positioned to detect conflicts in memory-driven representations arriving from CA3 and sensory information from entorhinal cortex, with new CA2 cell assemblies being recruited upon small environmental changes (Wintzer et al., 2014). According to this hypothesis, novelty induced activity in CA2 may forward sensory-driven information to downstream CA1 while inhibiting memory-driven output from CA3. Following this hypothesis CA2-silenced animals may lack the circuitry to quickly signal a change in the platform location and thus show more perseverance. Whether newly recruited CA2 assemblies are meant to store the memory directly or only relay the signal to adjacent hippocampal subregions is not yet clear.

In our recent theoretical work we postulate that CA2 prioritizes important experiences for replay during hippocampal sharp wave ripples (Stöber et al., 2020; Lehr et al., 2021). Neuromodulator release in CA2 due to novelty or saliency may unlock plasticity between CA3-CA2 and pair co-active assemblies in the two regions. This pairing of assembly sequences would then shift the probability distribution of replay to facilitate integrating new information into internal representations. Thus without CA2, animals would struggle to consolidate knowledge about the new platform location because neural activity sequences representing paths to the new location are not sufficiently prioritized over neural activity sequences related to the original platform. Given that hippocampal replay is also involved in planning (Pfeiffer and Foster, 2013), reduced prioritization of assembly sequences related to the relocated platform location could also disrupt planning-related replay events during the task. The effect on consolidation or planning could arise across trials during reversal learning, or beginning after the probe trial on day 8, where animals first experience that the original platform is removed.

Notably, both the memory/sensory switch and the prioritization hypothesis are complimentary. A change in platform location would certainly constitute a very important survival experience for an animal and would likely both induce neuromodulator release at CA3-CA2 synapses and recruit new CA2 assemblies. Paired CA3-CA2 sequences would then be prioritized for consolidation into memory due to neuromodulator-dependent pairing. Newly recruited CA2 neurons may support the encoding of paths to the new platform location or simply add a unique identifier to the existing hippocampal population code induced by the novelty of the situation.

### 4.3 Working memory deficits in CA2-silenced mice as alternative explanation for perseverance in reversal learning

A deficit in working memory in CA2-silenced mice could also contribute to slower reversal learning in the water task. Recent work has shown that silencing excitatory input from dorsal CA2 to dorsal CA1 during a 10 second delay period impairs performance on a delayed spatial alternation task (MacDonald and Tonegawa, 2021a, see also Lehr and Stöber, 2021; MacDonald and Tonegawa, 2021b). If working memory was disrupted in the current study, not remembering what happened in the preceding minutes could explain perseverance at the old platform location in subsequent trials. However if working memory were impaired, one might expect difficulties in both learning and reversal learning. Though it is conceivable that an effect of impaired working memory would be more prominent in reversal learning since animals may rely on the established memory of the previous platform location and have difficulties holding information about the new platform location in working memory from the most recent trial. It is worth mentioning that the working memory deficit in the literature (MacDonald and Tonegawa, 2021a) is after a 10 second delay period, while trial times in reversal learning in this study were separated by around 20 minutes.

### 4.4 CA2’s position in the DG → CA2 → vCA1 circuit may explain reduced flexibility

Another possible explanation for dorsal CA2 involvement in behavioral flexibility during reversal learning is its position in the DG → CA2 → vCA1 circuit. The DG projects to dorsal CA2 (Kohara et al., 2014) and newborn DG granule cells establish new synaptic contacts with CA2 pyramidal neurons (Llorens-Martín et al., 2015). If growth of new DG granule cells is suppressed, animals show increased perseverance at the old goal location after platform relocation in the water task, comparable to the CA2-silenced animals in this study (Garthe et al., 2009). The loss of new connections from newborn DG neurons to the dorsal CA2 could therefore account for increased perseverance in CA2-silenced animals.

Further, dorsal CA2 projects to the ventral hippocampus (Kohara et al., 2014; Meira et al., 2018; Okuyama et al., 2016; Tamamaki et al., 1988) which has been implicated in early learning in the Morris water task. Ruediger et al. (2012) showed that ventral hippocampus was needed for local search strategies (scanning and chaining) that are required early on during acquisition and lesions resulted in delayed onset of goal-directed spatial search strategies (directed search, focal search, and direct swim). Despite impairments in early learning, lesioned animals were still able to learn the platform location over multiple training days. Their pattern of results is very similar to ours: animals overcame a delay in the development of spatial search and had no impairments in the probe trial. In contrast to our study, Ruediger et al. (2012) described an involvement of ventral hippocampus during initial acquisition and did not investigate reversal learning.

Thus inactivation of dorsal CA2 projections to ventral hippocampus could also contribute to the deficits observed in CA2-silenced mice. The effect may be mediated by ventral CA1 projections to striatal reward circuitry (Ruediger et al., 2012). With dorsal CA2 silenced, the link between newborn DG granule cells and the ventral CA1 would be weakened, and as such, it is conceivable that important information would not get through the hippocampus to the striatum.

### 4.5 Dorsal CA2 supports flexibility in early reversal learning but was not required for initial learning

While we observed an effect of dorsal CA2 silencing on reversal learning, initial learning of the water task was not impaired. Thus we could not support the hypothesis that dorsal CA2 is recruited when forming a new representation, but instead seems to be recruited when representations need to be updated. This could, for example, stem from involvement in signalling a conflict in sensory inputs vs. memory content (Middleton and McHugh, 2019) or when novel memory traces need to be prioritized over old ones (Stöber et al., 2020). Whether disrupted working memory (MacDonald and Tonegawa, 2021a) or CA2’s position connecting dorsal dentate gyrus with ventral CA1 can explain the selective effect in reversal learning is less clear.

It does however remain possible that dorsal CA2 is recruited in initial learning but the procedure in this study was not able to detect the effects. Here, mice were housed in groups, habituated to handling and transport between the colony room and behavioral room for three days and then were exposed to two days of cued learning, which may have an influence on initial learning of the hidden platform on days 3 to 7. It could be that differences in CA2-silenced mice may be detectable by using a protocol like Ruediger and colleagues, in which animals were housed individually (see Loisy et al., this issue) and no cued phase preceded initial learning. Under those conditions, Ruediger et al. (2012) observed robust effects of subdivision-specific hippocampal lesions on initial learning. It is also worth considering that the transition from a cued platform to a hidden platform may require updating a representation, however in this case the task in both phases is not the same. During cued learning the platform was moved between every trial and so learning a spatial location is not required.

Nevertheless, the current data support the notion that dorsal CA2 is required to resolve conflicting information and update representations, not for formation of completely new ones. Interestingly, Ruediger et al. (2012) showed a progression of involvement during learning from ventral through intermediate to dorsal hippocampus over the course of nine days. It could be that early learning depends primarily on ventral hippocampus while early reversal learning may depend on the whole hippocampus, including dorsal CA2, to reconcile the new representation with the old.

### 4.6 Further experiments can elucidate the contribution of CA2 in spatial learning

To discern whether one or more of the hypotheses outlined above indeed account for CA2 involvement in behavioral flexibility in spatial learning, further experiments must be conducted. It is important to note, that despite perseverance for the old location during reversal learning, the animals were still able to acquire and remember the novel location. This indicates that five days of reversal learning is sufficient to compensate for the deleterious effects of more perseverance. It also suggests that some tasks might be better for detecting the type of subtle learning and memory impairments elicited by CA2 manipulations. A single massed training session to a new location and a probe trial 24 hours later (Bolding and Rudy, 2006; Deibel et al., 2014, 2022), would most likely reveal impairments in both learning and retention. Along these lines CA2-silenced mice would likely also be impaired at finding a new spatial location every training day (Whishaw, 1985). Further consideration should also be given to the effect of the probe trial before reversal learning, where animals have their first opportunity to learn that the platform is no longer available, as well as the time between trials. Both could influence whether consolidation or working memory is recruited by the task.

Additionally, it will be key to test the hippocampal response to conflicts in sensory vs. memory information in CA2-silenced and control animals with electrophysiological measurements across the hippocampus. According to the memory/sensory switch hypothesis, activity in CA3 should primarily carry information about the old, but not the new platform location in the first trials after encountering the new platform. In contrast, newly recruited cell assemblies in CA2 should carry information about the novel platform and contribute to a strong inhibition of CA3 (Middleton and McHugh, 2019), or alternatively, indirectly weaken CA3’s information flow to CA1.

If prioritized replay of salient experiences explains CA2 involvement, we would expect increased and coordinated activation of CA3 and CA2 assembly sequences representing trajectories to the novel platform location beginning after the first reversal learning trial, and likely changes in replay after the probe trial related to the removal of the original platform. Silencing CA2 during or after the trial or blocking CA2-specific neuromodulation during encoding should prevent prioritized reactivation. Such an experiment could also elucidate whether CA3-CA2 pairing only plays a role in consolidation or is also required in planning, as this should be observable in replay distributions during rest vs. during navigation in subsequent trials. If the effect is dependent on synaptic consolidation, which is recruited at CA3-CA2 synapses (cf. synaptic tagging and capture, Dasgupta et al., 2017, 2020; Benoy et al., 2018, 2022), then inhibiting protein synthesis in CA2 directly after reversal learning would block pairing of CA2 and CA3 assemblies and thus disrupt prioritization, thereby eliciting a similar effect to CA2 silencing.

It will be crucial to specifically test flexible reversal learning in transiently silenced CA2 animals, either selectively during the task or in the minutes to hours afterwards, with simultaneous electrophysiological recordings across the hippocampus. Selectively silencing CA2 during certain parts of the task will distinguish between involvement in encoding, consolidation, recall, and planning and thereby help elucidate the underlying neural mechanisms. Furthermore, transient silencing would also avoid the potential confounding factor that mice with chronically silenced CA2 have been shown to develop spatially triggered network hyperexcitability events, which are reminiscent of spike-wave-type discharges associated with seizures (Boehringer et al., 2017). These events could negatively affect performance in the Morris water maze in a similar way as induced seizures (Gilbert et al., 2000).

### 4.7 Conclusion

Here we report that silencing dorsal CA2 reduces behavioral flexibility during reversal learning in the Morris water task. This is important because it directly demonstrates that CA2 is fundamentally involved in hippocampus-dependent learning and memory. We provide testable predictions to dissect the specific mechanisms underlying CA2’s contribution to behavioral flexiblility.

## 5 Acknowledgements

We thank Rebecca Piskorowski and her team as well as Arvind Kumar for providing valuable feedback on the interpretation of these findings. We thank Steven Siegelbaum for sharing the data.

## 7 Data Availability

Data from (Hitti and Siegelbaum, 2014), stored in the expipe format (Lepperød et al., 2020), and code are available at https://github.com/tristanstoeber/CA2_spatiallearning.

## References

Alexander, G. M., Farris, S., Pirone, J. R., Zheng, C., Colgin, L. L., and Dudek, S. M. (2016). Social and novel contexts modify hippocampal CA2 representations of space. Nature Communications, 7(1):1–14.

Benoy, A., Dasgupta, A., and Sajikumar, S. (2018). Hippocampal area CA2: an emerging modulatory gateway in the hippocampal circuit. Experimental Brain Research, pages 1–13.

Benoy, A., Wong, L.-W., Ather, N., and Sajikumar, S. (2022). Serotonin facilitates late-associative plasticity via synaptic tagging/cross-tagging and capture at hippocampal ca2 synapses in male rats. Oxford Open Neuroscience, 1.

Berdugo-Vega, G., Arias-Gil, G., López-Fernández, A., Artegiani, B., Wasielewska, J. M., Lee, C.-C., Lippert, M. T., Kempermann, G., Takagaki, K., and Calegari, F. (2020). Increasing neurogenesis refines hippocampal activity rejuvenating navigational learning strategies and contextual memory throughout life. Nature communications, 11(1):1–12.

Berdugo-Vega, G., Lee, C.-C., Garthe, A., Kempermann, G., and Calegari, F. (2021). Adult-born neurons promote cognitive flexibility by improving memory precision and indexing. Hippocampus, 31(10):1068–1079.

Berger, S., Wolfer, D. P., Selbach, O., Alter, H., Erdmann, G., Reichardt, H. M., Chepkova, A. N., Welzl, H., Haas, H. L., Lipp, H.-P., et al. (2006). Loss of the limbic mineralocorticoid receptor impairs behavioral plasticity. Proceedings of the National Academy of Sciences, 103(1):195–200.

Boehringer, R., Polygalov, D., Huang, A. J., Middleton, S. J., Robert, V., Wintzer, M. E., Piskorowski, R. A., Chevaleyre, V., and McHugh, T. J. (2017). Chronic loss of CA2 transmission leads to hippocampal hyperexcitability. Neuron, 94(3):642–655.

Bolding, K. and Rudy, J. W. (2006). Place learning in the morris water task: Making the memory stick. Learning & Memory, 13(3):278–286.

Boyle, L., Posani, L., Irfan, S., Siegelbaum, S. A., and Fusi, S. (2022). The geometry of hippocampal CA2 representations enables abstract coding of social familiarity and identity. bioRxiv.

Chevaleyre, V. and Siegelbaum, S. A. (2010). Strong CA2 pyramidal neuron synapses define a powerful disynaptic cortico-hippocampal loop. Neuron, 66(4):560–572.

Cope, E. C., Zych, A. D., Katchur, N. J., Waters, R. C., Laham, B. J., Diethorn, E. J., Park, C. Y., Meara, W. R., and Gould, E. (2021). Atypical perineuronal nets in the ca2 region interfere with social memory in a mouse model of social dysfunction. Molecular Psychiatry, pages 1–12.

Dasgupta, A., Baby, N., Krishna, K., Hakim, M., Wong, Y. P., Behnisch, T., Soong, T. W., and Sajikumar, S. (2017). Substance P induces plasticity and synaptic tagging/capture in rat hippocampal area CA2. Proceedings of the National Academy of Sciences, 114(41):E8741–E8749.

Dasgupta, A., Lim, Y. J., Kumar, K., Baby, N., Pang, K. L. K., Benoy, A., Behnisch, T., and Sajikumar, S. (2020). Group III metabotropic glutamate receptors gate long-term potentiation and synaptic tagging/capture in rat hippocampal area CA2. eLife, 9:e55344.

Deibel, S. H., Ingram, M. L., Lehr, A. B., Martin, H. C., Skinner, D. M., Martin, G. M., Hughes, I. M., and Thorpe, C. M. (2014). In a daily time–place learning task, time is only used as a discriminative stimulus if each daily session is associated with a distinct spatial location. Learning & behavior, 42(3):246–255.

Deibel, S. H., Lewis, L. M., Cleary, J., Cassell, T. T., Skinner, D. M., and Thorpe, C. M. (2022). Unpredictable mealtimes rather than social jetlag affects acquisition and retention of hippocampal dependent memory. Behavioural Processes, 201:104704.

Dodero, L., Damiano, M., Galbusera, A., Bifone, A., Tsaftsaris, S. A., Scattoni, M. L., and Gozzi, A. (2013). Neuroimaging evidence of major morpho-anatomical and functional abnormalities in the btbr t+ tf/j mouse model of autism. PloS one, 8(10):e76655.

Donegan, M. L., Stefanini, F., Meira, T., Gordon, J. A., Fusi, S., and Siegelbaum, S. A. (2020). Coding of social novelty in the hippocampal CA2 region and its disruption and rescue in a 22q11.2 microdeletion mouse model. Nature Neuroscience, 23(11):1365–1375.

Dupret, D., O’Neill, J., Pleydell-Bouverie, B., and Csicsvari, J. (2010). The reorganization and reactivation of hippocampal maps predict spatial memory performance. Nature Neuroscience, 13(8):995–1002.

Ego-Stengel, V. and Wilson, M. A. (2010). Disruption of ripple-associated hippocampal activity during rest impairs spatial learning in the rat. Hippocampus, 20(1):1–10.

Ellegood, J. and Crawley, J. N. (2015). Behavioral and neuroanatomical phenotypes in mouse models of autism. Neurotherapeutics, 12(3):521–533.

Garthe, A., Behr, J., and Kempermann, G. (2009). Adult-generated hippocampal neurons allow the flexible use of spatially precise learning strategies. PloS one, 4(5):e5464.

Gilbert, T. H., Hannesson, D. K., and Corcoran, M. E. (2000). Hippocampal kindled seizures impair spatial cognition in the morris water maze. Epilepsy research, 38(2-3):115–125.

Girardeau, G., Benchenane, K., Wiener, S. I., Buzsáki, G., and Zugaro, M. B. (2009). Selective suppression of hippocampal ripples impairs spatial memory. Nature Neuroscience, 12(10):1222.

Gozzi, A. and Schwarz, A. J. (2016). Large-scale functional connectivity networks in the rodent brain. Neuroimage, 127:496–509.

He, H., Boehringer, R., Huang, A. J., Overton, E. T., Polygalov, D., Okanoya, K., and McHugh, T. J. (2021). CA2 inhibition reduces the precision of hippocampal assembly reactivation. Neuron, 109(22):3674–3687.

He, H., Wang, Y., and McHugh, T. J. (2022). Behavioral status modulates CA2 influence on hippocampal network dynamics. Hippocampus.

Hitti, F. L. and Siegelbaum, S. A. (2014). The hippocampal CA2 region is essential for social memory. Nature, 508(7494):88–92.

Jadhav, S. P., Kemere, C., German, P. W., and Frank, L. M. (2012). Awake hippocampal sharp-wave ripples support spatial memory. Science, 336(6087):1454–1458.

Kohara, K., Pignatelli, M., Rivest, A. J., Jung, H.-Y., Kitamura, T., Suh, J., Frank, D., Kajikawa, K., Mise, N., Obata, Y., et al. (2014). Cell type-specific genetic and optogenetic tools reveal hippocampal CA2 circuits. Nature Neuroscience, 17(2):269–279.

Kretz, O., Schmid, W., Berger, S., and Gass, P. (2001). The mineralocorticoid receptor expression in the mouse cns is conserved during development. Neuroreport, 12(6):1133–1137.

Lee, H., Wang, C., Deshmukh, S. S., and Knierim, J. J. (2015). Neural population evidence of functional hetero-geneity along the CA3 transverse axis: pattern completion versus pattern separation. Neuron, 87(5):1093–1105.

Lee, S. E., Simons, S. B., Heldt, S. A., Zhao, M., Schroeder, J. P., Vellano, C. P., Cowan, D. P., Ramineni, S., Yates, C. K., Feng, Y., et al. (2010). RGS14 is a natural suppressor of both synaptic plasticity in CA2 neurons and hippocampal-based learning and memory. Proceedings of the National Academy of Sciences, 107(39):16994–16998.

Lehr, A. B., Kumar, A., Tetzlaff, C., Hafting, T., Fyhn, M., and Stöber, T. M. (2021). CA2 beyond social mem-ory: Evidence for a fundamental role in hippocampal information processing. Neuroscience & Biobehavioral Reviews, 126:398–412.

Lehr, A. B. and Stöber, T. M. (2021). Differential involvement of ca2 in internally vs. externally driven hippocam-pal sequences. Proceedings of the National Academy of Sciences, 118(38):e2110671118.

Lepperød, M. E., Dragly, S.-A., Buccino, A. P., Mobarhan, M. H., Malthe-Sørenssen, A., Hafting, T., and Fyhn, M. (2020). Experimental pipeline (expipe): A lightweight data management platform to simplify the steps from experiment to data analysis. Frontiers in Neuroinformatics, 14:30.

Llorens-Martín, M., Jurado-Arjona, J., Avila, J., and Hernández, F. (2015). Novel connection between newborn granule neurons and the hippocampal CA2 field. Experimental neurology, 263:285–292.

MacDonald, C. J. and Tonegawa, S. (2021a). Crucial role for CA2 inputs in the sequential organization of CA1 time cells supporting memory. Proceedings of the National Academy of Sciences, 118(3).

MacDonald, C. J. and Tonegawa, S. (2021b). Reply to lehr and stöber: What’s in a name? on the distinction between temporal coding and internally driven activity. Proceedings of the National Academy of Sciences, 118(38):e2112026118.

McCann, K. E., Lustberg, D. J., Shaughnessy, E. K., Carstens, K. E., Farris, S., Alexander, G. M., Radzicki, D., Zhao, M., and Dudek, S. M. (2019). Novel role for mineralocorticoid receptors in control of a neuronal phenotype. Molecular psychiatry, pages 1–15.

Meira, T., Leroy, F., Buss, E. W., Oliva, A., Park, J., and Siegelbaum, S. A. (2018). A hippocampal circuit linking dorsal CA2 to ventral CA1 critical for social memory dynamics. Nature Communications, 9(1):4163.

Middleton, S. J. and McHugh, T. J. (2019). CA2: A highly connected intrahippocampal relay. Annual Review of Neuroscience, 43:2020.

Morris, R. (1984). Developments of a water-maze procedure for studying spatial learning in the rat. Journal of neuroscience methods, 11(1):47–60.

Morris, R. G., Garrud, P., Rawlins, J. a., and O’Keefe, J. (1982). Place navigation impaired in rats with hippocam-pal lesions. Nature, 297(5868):681–683.

Moy, S. S., Nadler, J. J., Young, N. B., Perez, A., Holloway, L. P., Barbaro, R. P., Barbaro, J. R., Wilson, L. M., Threadgill, D. W., Lauder, J. M., et al. (2007). Mouse behavioral tasks relevant to autism: phenotypes of 10 inbred strains. Behavioural brain research, 176(1):4–20.

Nakazawa, K., Sun, L. D., Quirk, M. C., Rondi-Reig, L., Wilson, M. A., and Tonegawa, S. (2003). Hippocampal ca3 nmda receptors are crucial for memory acquisition of one-time experience. Neuron, 38(2):305–315.

Okuyama, T., Kitamura, T., Roy, D. S., Itohara, S., and Tonegawa, S. (2016). Ventral CA1 neurons store social memory. Science, 353(6307):1536–1541.

Oliva, A., Fernández-Ruiz, A., Leroy, F., and Siegelbaum, S. A. (2020). Hippocampal CA2 sharp-wave ripples reactivate and promote social memory. Nature, 587(7833):264–269.

Overall, R. W., Zocher, S., Garthe, A., and Kempermann, G. (2020). Rtrack: a software package for reproducible automated water maze analysis. BioRxiv.

Pfeiffer, B. E. and Foster, D. J. (2013). Hippocampal place-cell sequences depict future paths to remembered goals. Nature, 497(7447):74–79.

R Core Team (2021). R: A Language and Environment for Statistical Computing. R Foundation for Statistical Computing, Vienna, Austria.

Ruediger, S., Spirig, D., Donato, F., and Caroni, P. (2012). Goal-oriented searching mediated by ventral hip-pocampus early in trial-and-error learning. Nature neuroscience, 15(11):1563–1571.

Singer, A. C., Carr, M. F., Karlsson, M. P., and Frank, L. M. (2013). Hippocampal swr activity predicts correct decisions during the initial learning of an alternation task. Neuron, 77(6):1163–1173.

Stöber, T. M., Lehr, A. B., Hafting, T., Kumar, A., and Fyhn, M. (2020). Selective neuromodulation and mutual inhibition within the CA3–CA2 system can prioritize sequences for replay. Hippocampus, 30(11):1228–1238.

Sutherland, R. J., Whishaw, I. Q., and Kolb, B. (1983). A behavioural analysis of spatial localization following electrolytic, kainate-or colchicine-induced damage to the hippocampal formation in the rat. Behavioural Brain Research, 7(2):133–153.

Tamamaki, N., Abe, K., and Nojyo, Y. (1988). Three-dimensional analysis of the whole axonal arbors originating from single CA2 pyramidal neurons in the rat hippocampus with the aid of a computer graphic technique. Brain Research, 452(1):255–272.

Tsien, J. Z., Huerta, P. T., and Tonegawa, S. (1996). The essential role of hippocampal CA1 nmda receptor– dependent synaptic plasticity in spatial memory. Cell, 87(7):1327–1338.

Whishaw, I. Q. (1985). Cholinergic receptor blockade in the rat impairs locale but not taxon strategies for place navigation in a swimming pool. Behavioral neuroscience, 99(5):979.

Wintzer, M. E., Boehringer, R., Polygalov, D., and McHugh, T. J. (2014). The hippocampal CA2 ensemble is sensitive to contextual change. The Journal of Neuroscience, 34(8):3056–3066.

Zhao, M., Choi, Y.-S., Obrietan, K., and Dudek, S. M. (2007). Synaptic plasticity (and the lack thereof) in hippocampal CA2 neurons. The Journal of Neuroscience, 27(44):12025–12032.

