## Supplemental information for "Silencing hippocampal CA2 reduces behavioral flexibility in spatial learning"

### Silencing hippocampal region CA2 leads to reduced flexibility in spatial learning

February 9, 2023

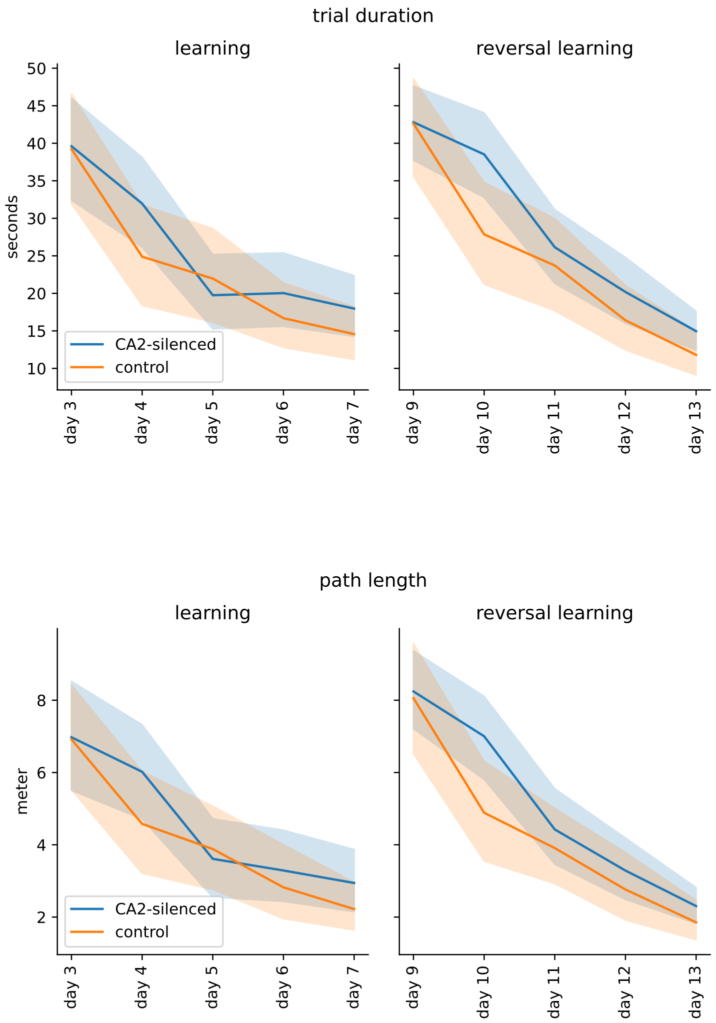


Supplementary Figure 1: **Trial duration and path length visually suggest subtle differences between CA2-silenced and control mice on certain days.** Reproduction of Fig 6b,c in Hitti & Siegelbaum (2014). Post-hoc tests with $p < 0.05$ for day 4, 10 and 13 for both measures (Mann-Whitney U test, one-tailed). However note two-way repeated measures ANOVA does not detect any significant treatment effect within the learning or reversal learning phase for either measure. Lines show the mean and shaded regions the $95\%$ confidence interval.

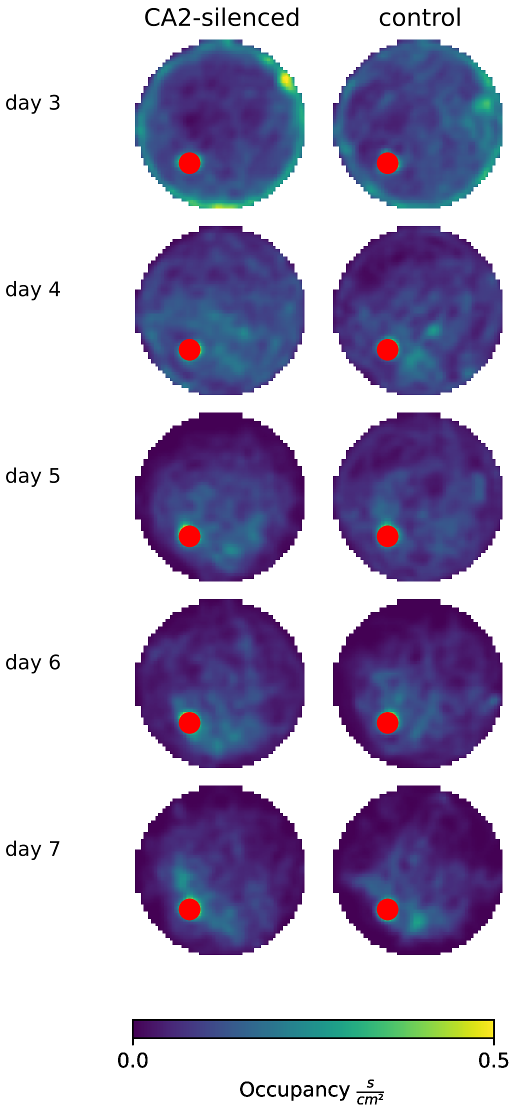


Supplementary Figure 2: **Average occupancy maps during the learning phase do not show a systematic difference between CA2-silenced and control mice.** Yellow represents high and blue low occupancy on a linear scale. Red circle demarcates the hidden platform location during learning.

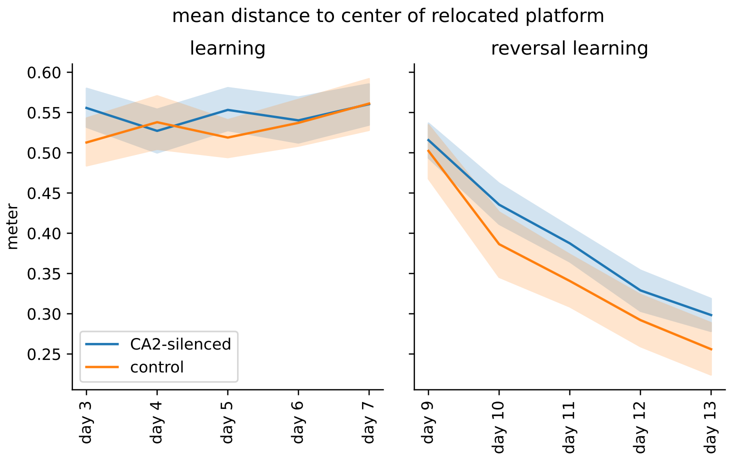


Supplementary Figure 3: **CA2-silenced mice trended towards a larger mean distance to the relocated platform location during reversal learning**. As expected, during initial learning the mean distance to the platform location that would be introduced in the reversal phase did not systematically differ between CA2-silenced animals and controls. However a consistent trend was observed during reversal learning (two-way repeated-measures ANOVA: $F(1, 16) = 2.92$, $p = 0.11$). Lines show the mean and shaded regions the $95\%$ confidence interval.
